# APOBEC1 mediated C-to-U RNA editing: target sequence and trans-acting factor contribution to 177 RNA editing events in 119 murine transcripts in-vivo

**DOI:** 10.1101/2021.01.08.425897

**Authors:** Saeed Soleymanjahi, Valerie Blanc, Nicholas O. Davidson

## Abstract

Mammalian C-to-U RNA editing was described more than 30 years ago as a single nucleotide modification in APOB RNA in small intestine, later shown to be mediated by the RNA-specific cytidine deaminase APOBEC1. Reports of other examples of C-to-U RNA editing, coupled with the advent of genome-wide transcriptome sequencing, identified an expanded range of APOBEC1 targets. Here we analyze the cis-acting regulatory components of verified murine C-to-U RNA editing targets, including nearest neighbor as well as flanking sequence requirements and folding predictions. We summarize findings demonstrating the relative importance of trans-acting factors (A1CF, RBM47) acting in concert with APOBEC1. Using this information, we developed a multivariable linear regression model to predict APOBEC1 dependent C-to-U RNA editing efficiency, incorporating factors independently associated with editing frequencies based on 103 Sanger-confirmed editing sites, which accounted for 84% of the observed variance. Co-factor dominance was associated with editing frequency, with RNAs targeted by both RBM47 and A1CF observed to be edited at a lower frequency than RBM47 dominant targets. The model also predicted a composite score for available human C-to-U RNA targets, which again correlated with editing frequency.

## INTRODUCTION

Mammalian C-to-U RNA editing was identified as the molecular basis for human intestinal APOB48 production more than three decades ago (Chen et al. 1987; Hospattankar et al. 1987; Powell et al. 1987). A site-specific enzymatic deamination of C6666 to U of *Apob* mRNA was originally considered the sole example of mammalian C-to-U RNA editing, occurring at a single nucleotide in a 14 kilobase transcript and mediated by an RNA specific cytidine deaminase (APOBEC1) (Teng et al. 1993). With the advent of massively parallel RNA sequencing technology we now appreciate that APOBEC1 mediated RNA editing targets hundreds of sites (Rosenberg et al. 2011; Blanc et al. 2014) mostly within 3’ untranslated regions of mRNA transcripts. This expanded range of targets of C-to-U RNA editing prompted us to reexamine key functional attributes in the regulatory motifs (both cis-acting elements and trans-acting factors) that impact editing frequency, focusing primarily on data emerging from studies of mouse cell and tissue-specific C-to-U RNA editing.

Earlier studies identified RNA motifs (Davies et al. 1989) contained within a 26-nucleotide segment flanking the edited cytidine base *in vivo* (in cell lines) or within 55 nucleotides using S100 extracts from rat hepatoma cells (Bostrom et al. 1989; Driscoll et al. 1989). Those, and other studies, established that *Apob* RNA editing reflects both the tissue/cell of origin as well as RNA elements remote and adjacent to the edited base (Bostrom et al. 1989; Davies et al. 1989). A granular examination of the regions flanking the edited base in *Apob* RNA demonstrated a critical 3’ sequence 6671-6681, downstream of C6666, in which mutations reduced or abolished editing activity (Shah et al. 1991). This 3’ site, termed a “mooring sequence” was associated with a 27s-“editosome” complex (Smith et al. 1991), which was both necessary and sufficient for site-specific *Apob* RNA editing and editosome assembly (Backus and Smith 1991). Other *cis*-acting elements include a 5 nucleotide spacer region between the edited cytidine and the mooring sequence, and also sequences 5’ of the editing site that regulate editing efficiency (Backus and Smith 1992; Backus et al. 1994) along with AU-rich regions both 5’ and 3’ of the edited cytidine that together function in concert with the mooring sequence (Hersberger and Innerarity 1998).

Advances in our understanding of physiological *Apob* RNA editing emerged in parallel from both the delineation of key RNA regions (summarized above) and also with the identification of components of the *Apob* RNA editosome (Sowden et al. 1996). APOBEC1, the catalytic deaminase (Teng et al. 1993) is necessary for physiological C-to-U RNA editing *in vivo* (Hirano et al. 1996) and *in vitro* (Giannoni et al. 1994). Using the mooring sequence of *Apob* RNA as bait, two groups identified APOBEC1 complementation factor (A1CF), an RNA-binding protein sufficient *in vitro* to support efficient editing in presence of APOBEC1 and *Apob* mRNA (Lellek et al. 2000; Mehta et al. 2000). Those findings reinforced the importance of both the mooring sequence and an RNA binding component of the editosome in promoting *Apob* RNA editing. However, while A1CF and APOBEC1 are sufficient to support *in vitro Apob* RNA editing, neither heterozygous (Blanc et al. 2005) or homozygous genetic deletion of *A1cf* impaired *Apob* RNA editing *in vivo* in mouse tissues (Snyder et al. 2017), suggesting that an alternate complementation factor was likely involved. Other work identified a homologous RNA binding protein, RBM47, that functioned to promote *Apob* RNA editing both *in vivo* and *in vitro* (Fossat et al. 2014), and more recent studies utilizing conditional, tissue-specific deletion of *A1cf* and *Rbm47* indicate that both factors play distinctive roles in APOBEC1-mediated C-to-U RNA editing, including *Apob* as well as a range of other APOBEC1 targets (Blanc et al. 2019).

These findings together establish important regulatory roles for both *cis*-acting elements and *trans*-acting factors in C-to-U mRNA editing. However, the majority of studies delineating *cis*-acting elements reflect earlier, *in vitro* experiments using *ApoB* mRNA and relatively little is known regarding the role of *cis*-acting elements in tissue-specific C-to-U RNA editing of other transcripts, *in vivo*. Here we use statistical modeling to investigate the independent roles of candidate regulatory factors in mouse C-to-U mRNA editing using data from *in vivo* studies from over 170 editing sites in 119 transcripts (Meier et al. 2005; Rosenberg et al. 2011; Gu et al. 2012; Blanc et al. 2014; Rayon-Estrada et al. 2017; Snyder et al. 2017; Blanc et al. 2019; Kanata et al. 2019). We also examined these regulatory factors in known human mRNA targets (Chen et al. 1987; Powell et al. 1987; Skuse et al. 1996; Mukhopadhyay et al. 2002; Grohmann et al. 2010; Schaefermeier and Heinze 2017).

## RESULTS

### Descriptive data

177 C-to-U RNA editing sites were identified based on eight studies that met inclusion and exclusion criteria (Meier et al. 2005; Rosenberg et al. 2011; Gu et al. 2012; Blanc et al. 2014; Rayon-Estrada et al. 2017; Snyder et al. 2017; Blanc et al. 2019; Kanata et al. 2019), representing 119 distinct RNA editing targets. 84% (100/119) of RNA targets were edited at one chromosomal location (Figure 1C) and 75% (89/119) of mRNA targets were edited at both a single chromosomal location and also within a single tissue (Figure 1D). The majority of editing sites occur in the 3’ untranslated region (142/177; 80%), with exonic editing sites the next most abundant subgroup (28/177; 16%, Figure 1E). Chromosome X harbors the highest number of editing sites (18/177; 10%), followed by chromosomes 2 and 3 (15/177; 8.5% for both, Supplemental Figure 1). 103/177 editing sites were confirmed by Sanger sequencing, with a mean editing frequency of 37 ± 22%.

**Figure 1.**
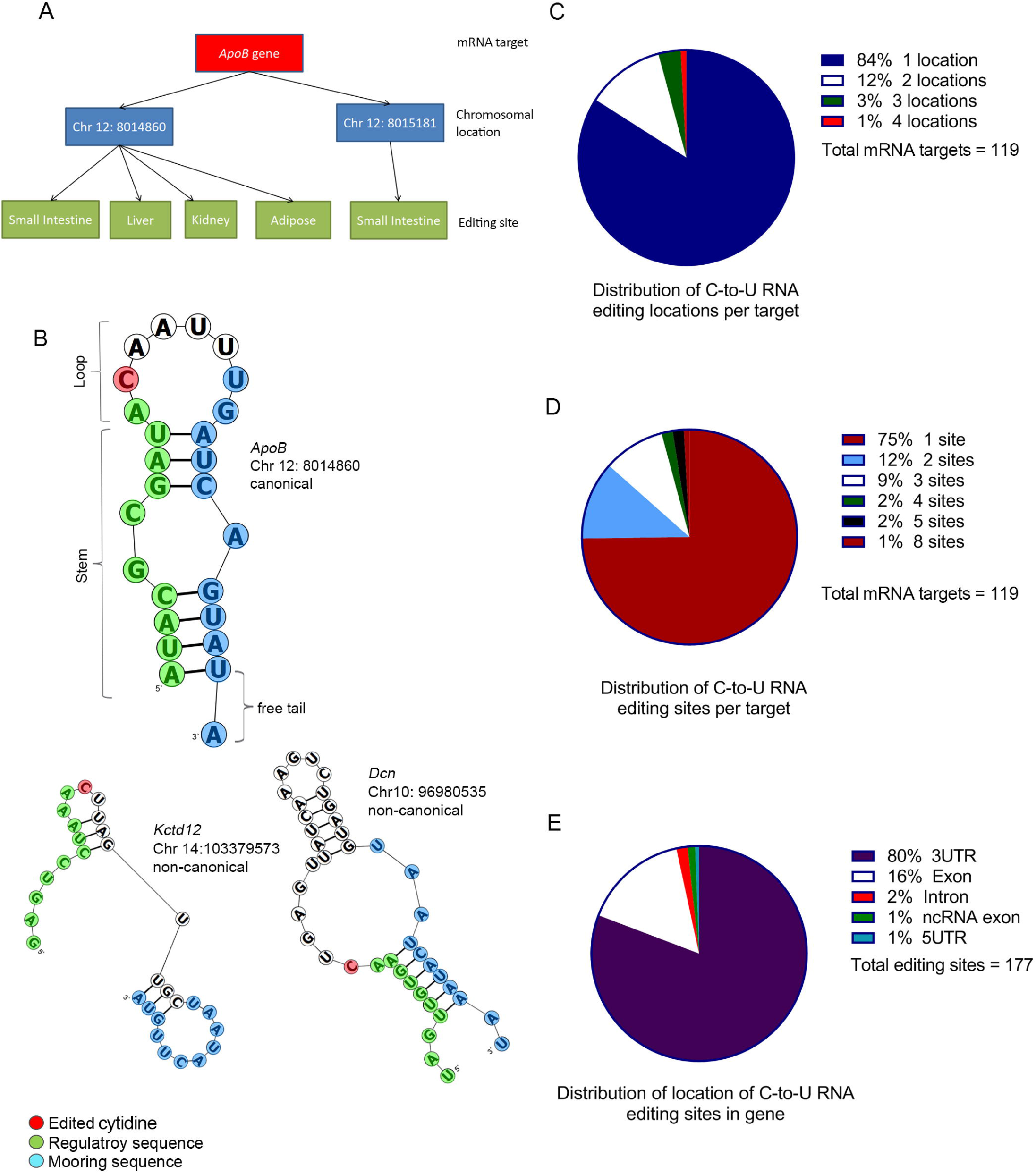
Characteristics of murine APOBEC1-mediated C-to-U mRNA editing sites. A: schematic presentation of mRNA target, chromosomal editing location, and editing sites considered. Each mRNA target could be edited at one or more chromosomal location(s) (blue boxes). Each editing location could be edited in one or more tissues giving rise to one or more editing site(s) per location (green boxes). Editing site(s) of each mRNA target are the sum of editing sites from all editing locations reported for that target. B: examples of canonical (*ApoB* chr12: 8014860, top) and two types of non-canonical (*Kctd12* chr14: 103379573 and *Dcn* chr10: 96980535) secondary structures. C: distribution of number of chromosomal editing location(s), or targeted cytidine(s), per mRNA target. D: distribution of number of total editing sites per mRNA target considering all chromosomal location(s) edited at different tissue(s). E: distribution of location of editing sites within gene structure.

### Base content of sequences flanking edited and mutated cytidines

AU content was enriched (∼87%) in nucleotides both immediately upstream and downstream of the edited cytidine across mouse RNA editing targets (Figure 2A and 2C). The average AU content across the region 10 nucleotides upstream to 20 nucleotides downstream of the edited cytidine was ∼70% (60 − 87%). Because APOBEC1 has been shown to be a DNA mutator (Harris et al. 2003; Wolfe et al. 2019; Wolfe et al. 2020), we determined the AU content of the mutated deoxycytidine region flanking human DNA targets (Nik-Zainal et al. 2012) to be ∼66% at a site one nucleotide downstream of the edited base (Figure 2B, C). The average AU content in the sequence 10 nucleotides upstream and 10 nucleotides downstream of mutated deoxycytidines is 59% (57-66.0%). The average AU content was 90% and 80% in nucleotides immediately upstream and downstream, respectively, of the targeted deoxycytidine in a subgroup of over 700 DNA editing events of the C to T type (Nik-Zainal et al. 2012), which is closer to the distribution found in C to U RNA editing targets. These features suggest that AU enrichment is an important component to editing function of APOBEC1 on both RNA and DNA targets, especially for the C/dC to U/dT change.

**Figure 2.**
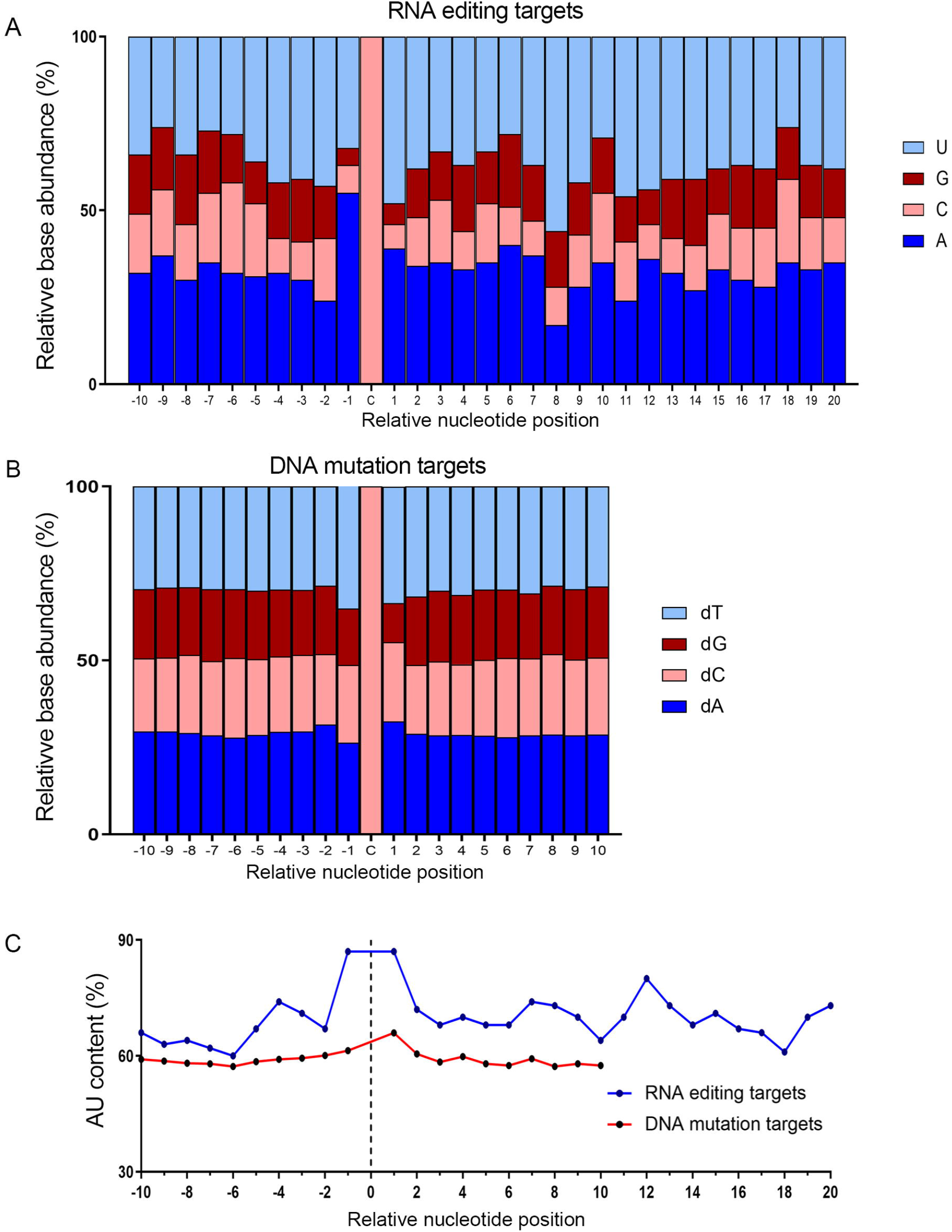
Base content of sequences flanking modified cytidine in RNA editing and DNA mutation targets. A: base content of 10 nucleotides upstream and 20 nucleotides downstream of edited cytidine in mouse APOBEC1-mediated C-to-U mRNA editing targets. B: base content of 10 nucleotides upstream and 10 nucleotides downstream of mutated cytidine in proposed human APOBEC-mediated DNA mutation targets in patients with breast cancer. C: comparison of AU base content (%) of nucleotides flanking modified cytidine in RNA editing targets and DNA mutation targets in mouse and human breast cancer patients, respectively.

### Factors influencing editing frequency

#### Regulatory-spacer-mooring cassette

We observed no significant associations between editing frequency and mismatches in motif A (*r*=-0.05, *P*=.46) or motif B (*r*=-0.1, *P*=.20) (Supplemental Figure 2), while mismatches in motif C and D negatively impacted editing frequency (*r*=-0.24, *P*=.001) (motif D *r*=-0.20, *P*=.008, Figure 3B). AU content of motif B showed a trend towards negative association with editing frequency (*r*=-0.13, *P*=.08 Figure 3C), but AU contents of motifs A (*r*=0.06, *P*=.4), C (*r*=-0.02, *P*=.8), and D (*r*=-0.02, *P*=.78) did not impact editing frequency (Supplemental Figure 2). The abundance of G in motif C (*r*=0.17, *P*=.02), abundance of C in motif B (*r*=0.13, *P*=.08), and G/C fraction in motif C (*r*=0.14, *P*=.04) showed either significance or a trend to associations with editing frequency. The spacer sequence averaged 5 ± 4 nucleotides, ranging from 0 to 20, with trend of association between length and editing frequency (*r*=-0.14, *P*=.09). The mean spacer sequence AU content was 73 ± 23%, with no association between editing frequency and AU content (*r*=-0.1, *P*=.2, Supplemental Figure 3). However, G abundance (*r*=-0.23, *P*=.01) and G/C fraction (*r*=-0.20, *P*=.03) of spacer showed significant associations with editing frequency in Sanger-confirmed targets. The mean number of mismatches in the first 4 nucleotides of the spacer sequence was 2.5 ± 1 with higher number of mismatches exerting a significant negative impact on editing frequency (*r*=-0.24, *P*=.01) (Figure 3D). The mean number of mismatches in the mooring sequence was 2.1 ± 1.8, ranging from 0 to 8 nucleotides. The number of mismatches showed a significant negative association with editing frequency (*r*=-0.30, *P*=.0003, Figure 3E). The base content of individual nucleotides surrounding the edited cytidine showed significant associations with editing frequency, which was more emphasized in nucleotides closer to the edited cytidine (Figure 3F, Supplemental Table 1). Furthermore, overall AU content of downstream sequence +16 to +20 had positive impact on editing frequency (*r*=0.17, *P*=.02) (Supplemental Figure 3). However, G abundance in downstream 20 nucleotides (*r*=-0.24, *P*=.001) and G/C fraction in downstream 10 nucleotides (*r*=-0.16, *P*=.09) showed significant or a trend of significant negative associations with editing frequency in Sanger-confirmed targets.

**Figure 3.**
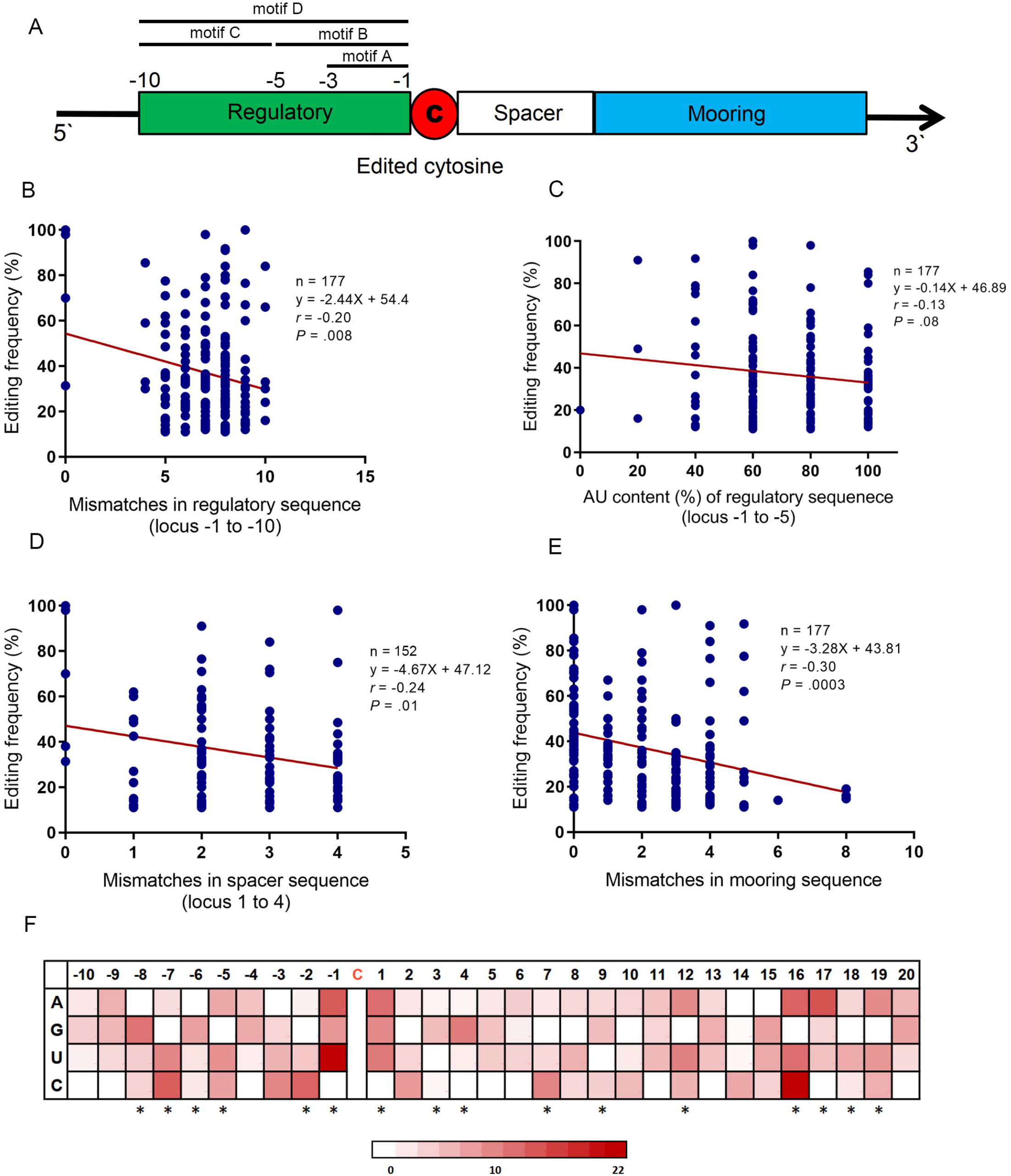
Characteristics of regulatory-spacer-mooring cassette and base content of individual nucleotides flanking edited cytidine in association with editing frequency. A: schematic illustration of regulatory-spacer-mooring cassette. Four motifs were defined for regulatory sequence: motif A for nucleotides -1 to -3; motif B for nucleotides -1 to -5; motif C for nucleotides -6 to -10; motif D representative of the whole sequence. B: association of the mismatches in motif D of regulatory sequence with editing frequency. C: association between the AU content (%) of regulatory sequence (motif B) and editing frequency. D: association of the mismatches in spacer (nucleotides +1 to +4 downstream of the edited cytidine) with editing frequency. E: association of the mismatches in mooring sequence with editing frequency. F: heatmap plot illustrating the association between base content of 30 nucleotides flanking the edited cytidine with editing frequency. Red color density in each cell represents the beta coefficient value of corresponding base in the multivariable linear regression model fit including that nucleotide. The asteriska refer to the nucleotides that were retained in the final model. Mismatches in regulatory, spacer, and mooring sequences were determined in comparison to the corresponding sequences in *ApoB* mRNA (as reference). *r*: Pearson correlation coefficient.

#### Secondary structure

We generated a predicted secondary structure for 172 editing sites, with four subgroups based on overall structure and location of the edited cytidine: loop (C_loop_), stem (C_stem_), tail (C_tail_), and non-canonical structure (NC). The majority of editing sites were in the C_loop_ subgroup (59%), followed by C_stem_ (20%), C_tail_ (13%), and NC (8%) subgroups (Figure 4A). Editing sites in the C_tail_ subgroup exhibited lower editing frequencies compared to editing sites in C_loop_ (29 ± 12 vs 41 ± 23%, *P*=.02) or C_stem_ (37 ± 21%, *P*=.04) subgroups. No significant differences were detected in other comparisons (Figure 4B). The edited cytidine was located in loop, stem, and tail of the secondary structure in 110 (64%), 38 (22%), and 24 (14%) of the edited RNAs, respectively. Editing sites with the edited cytidine within the loop exhibited significantly higher editing frequency compared to those with the edited cytidine in the tail (40 ± 24% vs 28 ± 12 %, *P*=.04). Other subgroups exhibited comparable editing frequencies (Supplemental Figure 4). The majority (78%) of editing sites contained a mooring sequence located in main stem-loop structure (Figure 4C), with the remainder located in the tail or secondary loop. Average editing efficiency was significantly higher in targets where the mooring sequence was located in the main stem-loop (Figure 4D). We also calculated the proportion of total nucleotides that constitute the main stem-loop in the secondary structure. The average ratio was 0.62 ± 0.18 ranging from 0.28 to 1 (Supplemental Table 2) with higher ratios associated with higher editing frequency of the corresponding editing site (*r*=0.20, *P*=.007) (Figure 4E). Finally, we considered the orientation of free tails in the secondary structure in terms of length and symmetry. Symmetric free tails were observed in 59% of editing sites (Supplemental Figure 4). The length of 5’ free tail showed negative association with editing frequency (*r*=-0.14, *P*=.04, Figure 4F) while no significant associations were detected between either the length of 3’ tail or symmetry of tails and editing frequency (Supplemental Figure 4).

**Figure 4.**
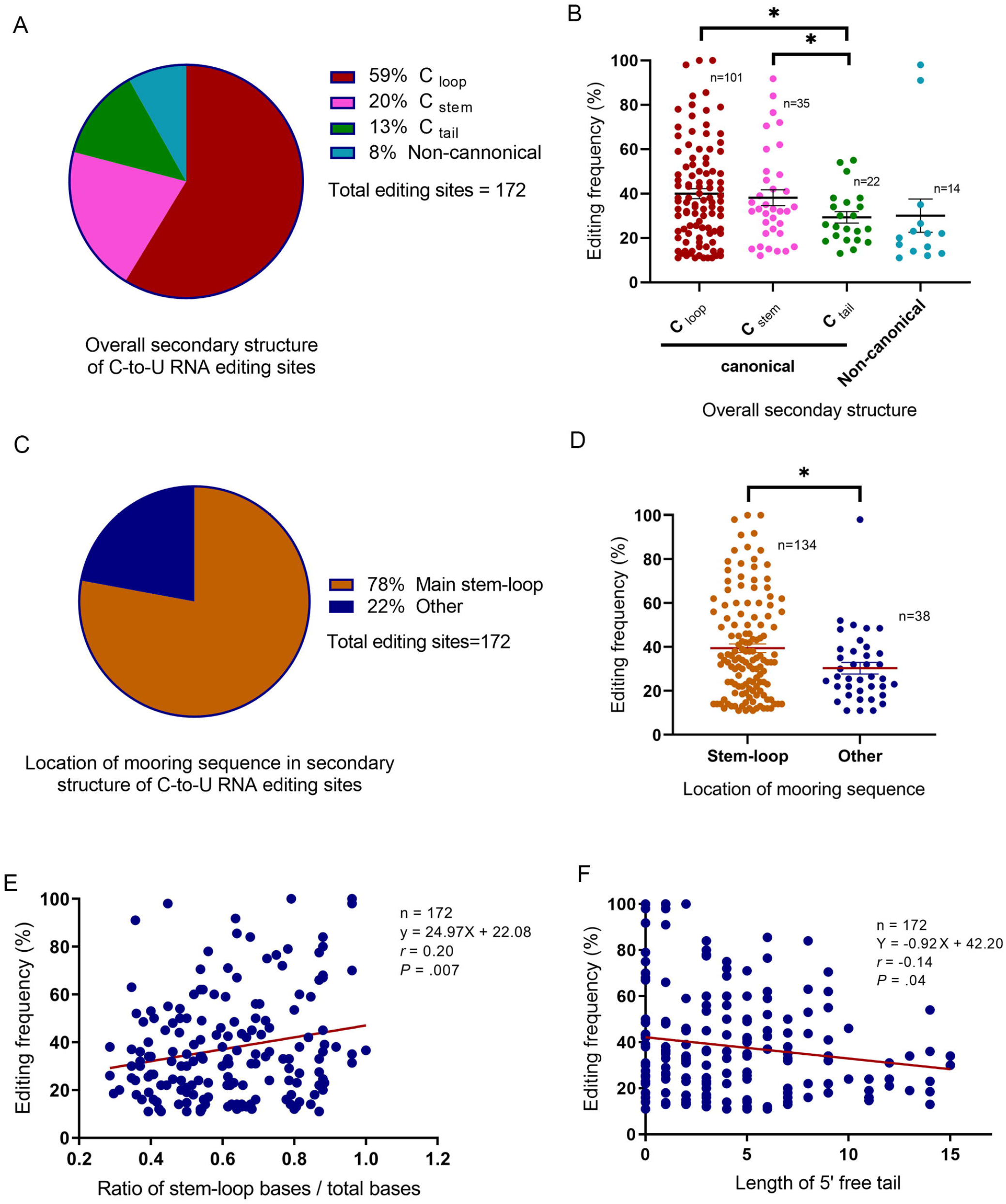
Secondary structure-related features in association with editing frequency. A: distribution of different types of overall secondary structure in editing sites. C _loop,_ C _stem,_ C _tail_ are three subtypes of canonical secondary structure based on the location of the edited cytidine. B: association between type of secondary structure and editing frequency. C: distribution of the mooring sequence location in editing sites. “Other” refers to mooring sequences located in tail or stem/loop and not part of the main stem-loop structure. D: association of mooring sequence location with editing frequency. E: association between ratio of main stem-loop bases to total bases count and editing frequency. F: association of the 5’ free tail length with editing frequency. * *P*<.05; ** *P*<.001. *r*: Pearson correlation coefficient.

#### *Trans*-acting factors and tissue specificity

Data for relative dominance of cofactors in APOBEC1-dependent RNA editing were available for 72 editing sites for targets in small intestine or liver (Blanc et al. 2019). RBM47 was identified as the dominant factor in 60/72 (83%) sites; A1CF was the dominant factor in 5/72 (7%) editing sites with the remaining sites (7/72; 10%), exhibiting equal codominancy (Figure 5A). The average editing frequencies at editing sites revealed differences across the groups with 41 ± 20% in RBM47-dominant targets, 23 ± 14% in A1CF-dominant, and 27 ± 11% in the co-dominant group (*P*=.03) (Figure 5B). The majority of RNA editing targets were edited in one tissue (103/119; 86% Figure 5C), while the maximum number of tissues in which an editing target is edited (at the same site) is 5 (*Cd36*). The small intestine harbors the highest number of verified editing sites (95/177; 54%), followed by liver (31/177; 17%), and adipose tissue (19/177; 11% Figure 5D). Sites edited in brain tissue showed the highest average editing frequency (54 ± 35 %, n=11), followed by bone marrow myeloid cells (50 ± 22 %, n=4), and kidney (47 ± 29%, n=10 Figure 5E).

**Figure 5.**
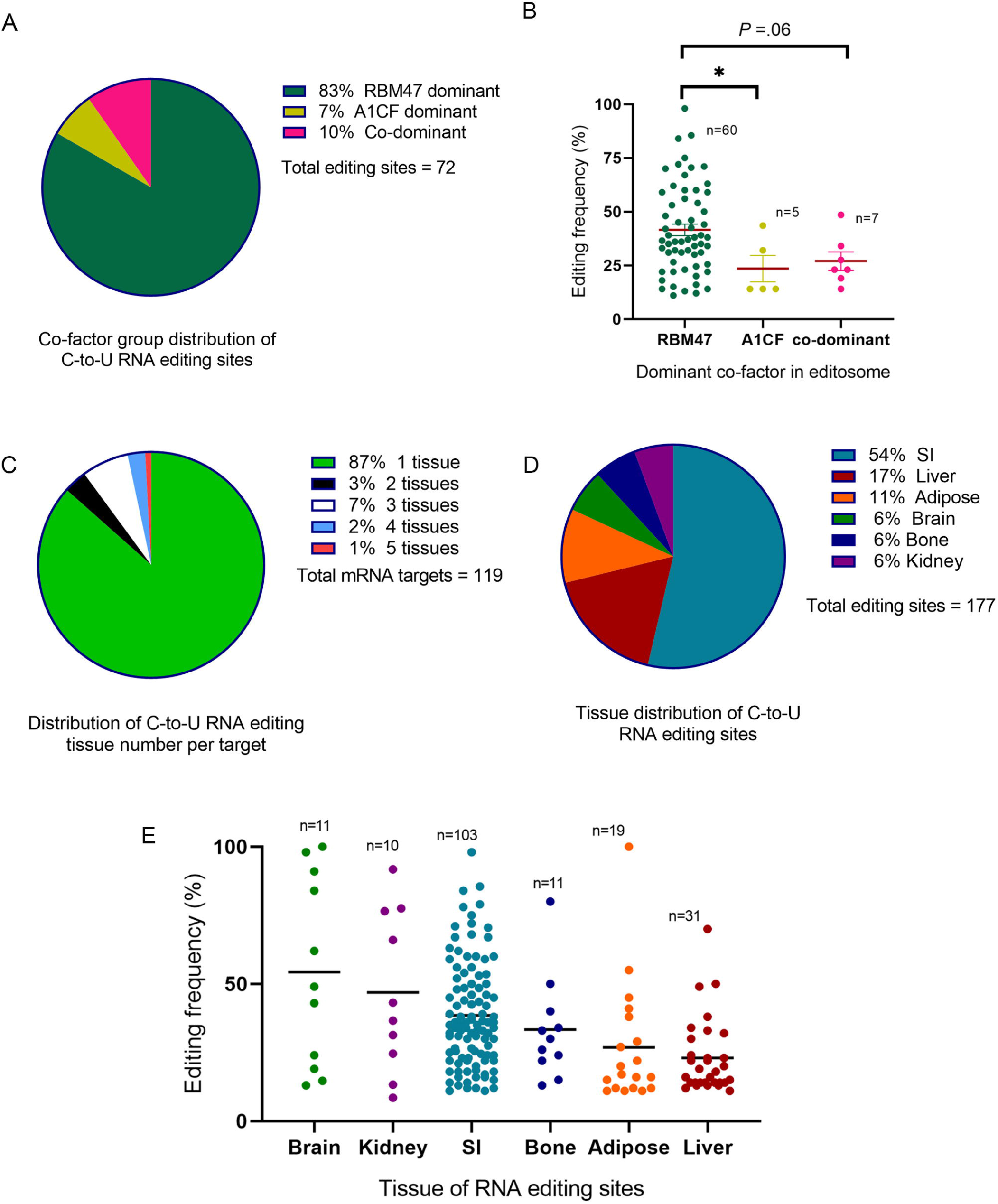
Dominance and tissue-specific cofactor patterns among editing sites. A: distribution of dominant co-factor in editosomes of editing sites. B: association of dominant co-factor with editing frequency. C: distribution of number of editing tissue(s) per mRNA target. D: tissue distribution of editing sites. E: average editing frequency of editing sites edited at different tissues. SI, small intestine.

We then developed a multivariable linear regression model to predict APOBEC1 dependent C-to-U RNA editing efficiency, incorporating factors independently associated with editing frequencies (Table 1). This model, based on 103 Sanger-confirmed editing sites with available data for all of the parameters mentioned, accounted for 84% of variance in editing frequency of editing sites included (R^2^=0.84, *P*<.001 Table 1). The final multivariable model revealed several factors independently associated with editing frequency, specifically the number of mismatches in mooring sequence; regulatory sequence motif D; AU content of regulatory sequence motif B; overall secondary structure for group C_tail_ vs group C_loop_; location of mooring sequence in secondary structure; “base content score” parameter that represents base content of the sequences flanking edited cytidine (Table 1). Removing “base content score” from the model reduced the power from R^2^=0.84 to R^2^=0.59. Next, we added a co-factor dominance variable and fit the model using the 72 editing sites with available data for cofactor dominance. Along with other factors mentioned above, co-factor dominance showed significant association with editing frequency (Table 1) with RNAs targeted by both RBM47 and A1CF observed to be edited at a lower frequency than RBM47 dominant targets.

**Table 1.**
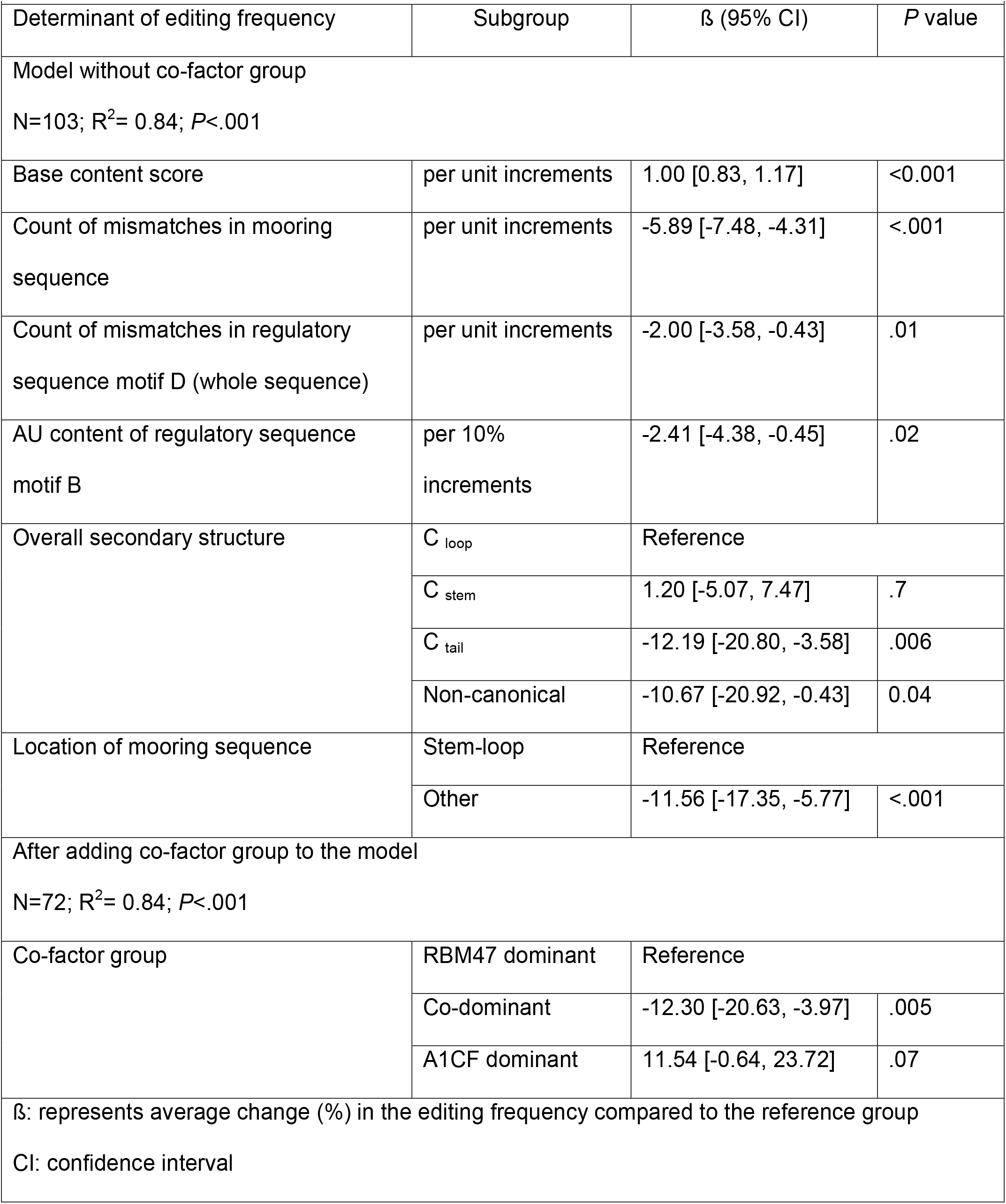
Multivariable linear regression model for determinant factors of editing frequency in mouse APOBEC1-dependent C-to-U mRNA editing sites.

Factors associated with co-factor dominance (Figure 6, Supplemental Table 3, Supplemental Figure 5), included tissue-specificity, with higher frequency of RBM47-dominant sites in small intestine compared to liver (91 vs 63%, *P*=.008) and A1CF-dominant and co-dominant editing sites more prevalent in liver. The number of mooring sequence mismatches also varied among three subgroups: 1.1 ± 1.3 in RBM47-dominant subgroup; 2.0 ± 2.5 in A1CF-dominant subgroup; and 2.9 ± 0.4 in co-dominant subgroup (*P*=.004). This was also the case regarding mismatches in the spacer: 2.4 ± 1.2 in RBM47-dominant subgroup; 2.7 ± 1.5 in A1CF-dominat subgroup; 3.8 ± 0.4 in co-dominant subgroup (*P*=.02). AU content (%) of downstream sequence +6 to +10 was higher in RBM47-dominant subgroup (P=.01). Finally, the location of the edited cytidine in secondary structure of mRNA strand was different across three subgroups (*P*=.04, Figure 6). We used pairwise multinomial logistic regression to determine factors independently associated with co-factor dominance (Figure 6C, Supplemental Table 4). C_tail_ editing sites, those with more mismatches in mooring and regulatory motif C, lower AU content in downstream sequence, and higher AU content in regulatory motif D were more likely co-dominant. Editing sites from small intestine and those with higher AU content of downstream sequence were more likely RBM47-dominant. Editing sites from liver and those with higher mismatches in regulatory motif B were more likely A1CF-dominant (Figure 6C).

**Figure 6.**
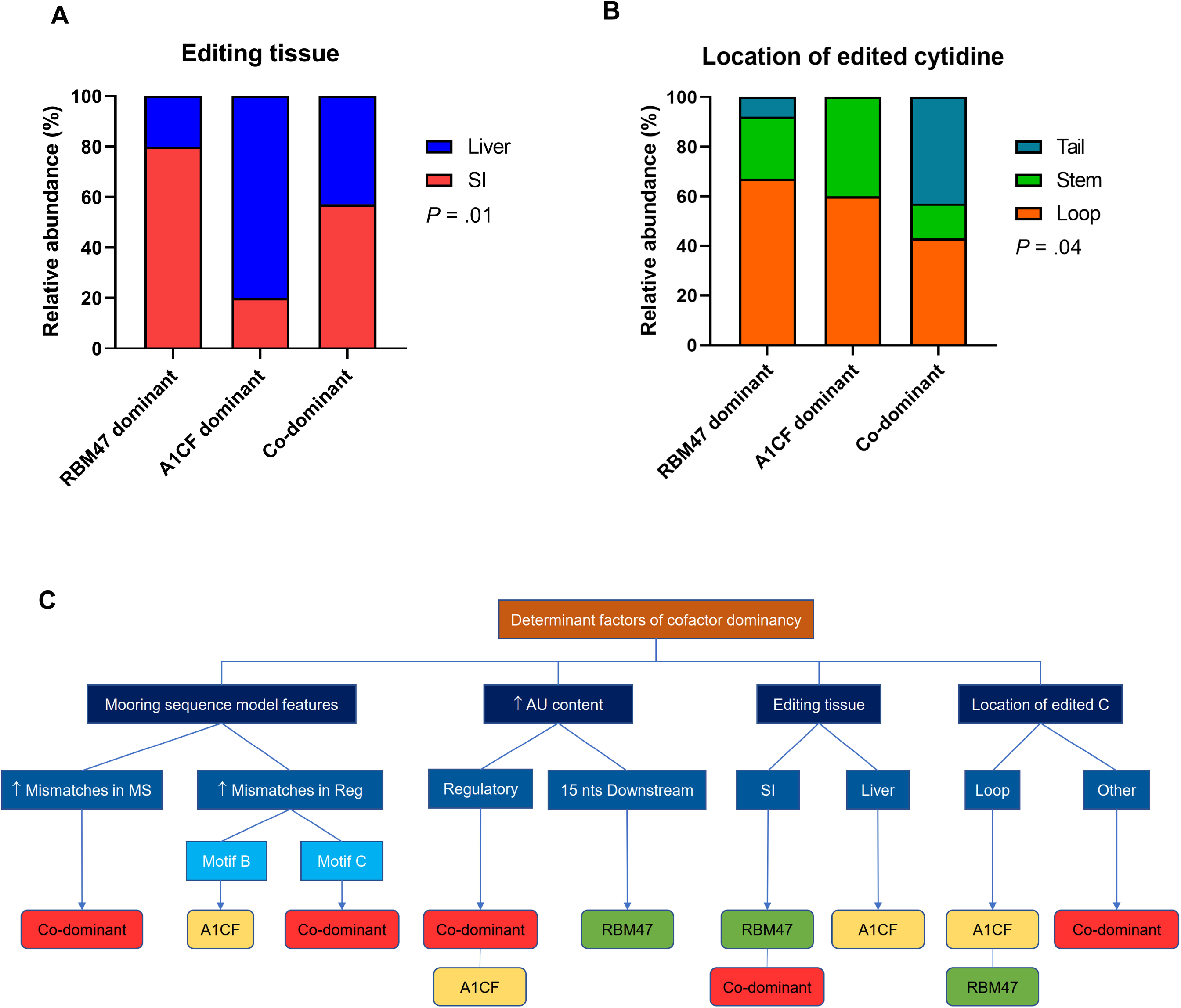
Co-factor pattern and tissue-specific role in murine C-to-U mRNA editing sites. A: distribution of editing tissue across subgroups of editing sites with different dominant co-factor patterns. B: location of edited cytidine in secondary structure of editing sites with different dominant co-factor patterns. C: schematic presentation of factors that correlate with dominant co-factor pattern in editing sites. This graph is based on the findings derived from pairwise multinomial logistic regression models.

### Human mRNA targets

Finally, we turned to an analysis of human C-to-U RNA editing targets for which this same panel of parameters was available (Table 2). Aside from *APOB* RNA, which is known to be edited in the small intestine (Chen et al. 1987; Powell et al. 1987), other targets have been identified in central or peripheral nervous tissue (Skuse et al. 1996; Mukhopadhyay et al. 2002; Meier et al. 2005; Schaefermeier and Heinze 2017). The human targets were categorized into low editing (*NF1, GLYR*_α_*2, GLYR*_α_*3*) and high editing (*APOB, TPH2B* exon3, *TPH2B* exon7) subgroups using 20% as cut-off. A composite score (maximum=6) was generated based on six parameters introduced in the mouse model with notable variance between the two subgroups including mismatches in mooring sequence, spacer length, location of the edited cytidine, and relative abundance of stem-loop bases (Table 2). High editing targets exhibited a significantly higher composite score (4.7 vs 2, *P*=.001) compared to low editing targets and the composite score significantly correlated with editing frequency in individual targets (*r*=0.95, *P*=.005). The canonical editing target ApoB (Chen et al. 1987; Powell et al. 1987) achieved a score of 5 (out of 6), reflecting the observation that one of the six parameters (AU% of regulatory motifs) in human *APOB* is non-preferential compared to the editing-promoting features identified in the mouse multivariable model.

**Table 2:**
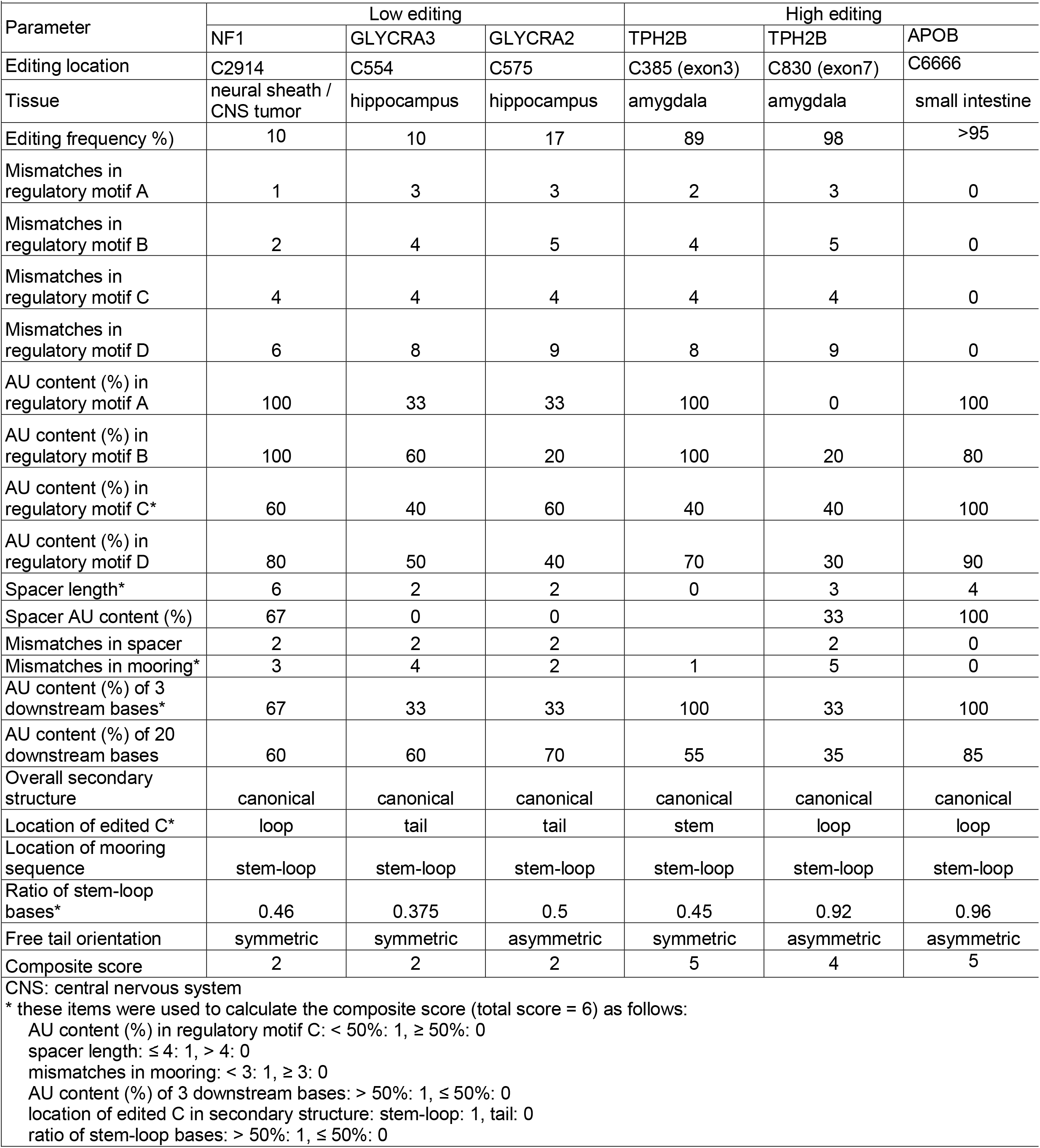
Characteristics of human C-to-U mRNA editing targets

## DISCUSSION

The current study reflects our analysis of 177 C-to-U RNA editing sites from 119 target mRNAs, with the majority residing within the 3’ untranslated region. Our multivariable model identified several key factors influencing editing frequency, including host tissue, base content of nucleotides surrounding the edited cytidine, number of mismatches in regulatory and mooring sequences, AU content of the regulatory sequence, overall secondary structure, location of the mooring sequence, and co-factor dominance. These factors, each exerting independent effects, together accounted for 84% of the variance in editing frequency. Our findings also showed that mismatches in the mooring and regulatory sequences, AU content of regulatory and downstream sequences, host tissue and secondary structure of target mRNA were associated with the pattern of co-factor dominance. Several aspects of these primary conclusions merit further discussion.

Previous studies investigating the key factors that regulate C-to-U mRNA editing were confined to *in vitro* studies and predicated on a single mRNA target (*ApoB*) (Backus and Smith 1991; Shah et al. 1991; Smith et al. 1991; Backus and Smith 1992; Hersberger and Innerarity 1998). With the expanded range of verified C-to-U RNA editing targets now available for interrogation, we revisited the original assumptions to understand more globally the determinants of C-to-U mRNA editing efficiency. In undertaking this analysis, we were reminded that the requirements for C-to-U mRNA editing *in vitro* often appear more stringent than *in vivo* (Backus and Smith 1991; Shah et al. 1991), which further emphasizes the importance of our findings. In addition, our approach included both *cis-acting* sequence- and folding-related predictions along with the role of *trans*-acting factors and took advantage of statistical modeling to adjust for confounding or modifier effects between these factors to identify their role in editing frequency.

We began with the assumptions established for *Apob* RNA editing which identified a 26 nucleotide segment encompassing the edited base, spacer, mooring sequence, and part of regulatory sequence as the minimal sequence competent for physiological editing *in vitro* and *in vivo* (Davies et al. 1989; Shah et al. 1991; Backus and Smith 1992). Those studies identified an 11-nucleotide mooring sequence as essential and sufficient for editosome assembly and site-specific C-to-U editing (Backus and Smith 1991; Shah et al. 1991; Backus and Smith 1992) and established optimal positioning of the mooring sequence relative to the edited base in *Apob* RNA (Backus and Smith 1992). The current work supports the key conclusions of this original mooring sequence model as applied to the entire range of C-to-U RNA editing targets. We observed that mismatches in either the mooring or regulatory sequences were independent factors governing editing frequency. By contrast, while mismatches in the spacer sequence also showed negative association with editing frequency, the impact of spacer mismatches were not retained in the final model, nor was the length of the spacer associated with editing frequency. Furthermore, we found mismatches in the regulatory sequence motif C to be more important than mismatches in motif B. These inconsistencies might conceivably reflect the context in which an RNA segment is studied (Backus and Smith 1992). For example, our analysis reflects physiological conditions in which naturally occurring mRNA targets are edited, while the aforementioned study used *in vitro* data based on varying lengths of *Apob* mRNA embedded within different mRNA contexts (*Apoe* RNA) (Backus and Smith 1992).

In addition to the components of mooring sequence model, we examined variations in the base content in different segments/motifs as well as among individual nucleotides surrounding the edited cytidine. As expected, we found that sequences flanking the edited cytidine exhibited high AU content. We further observed a similarly high AU content in the flanking sequences of a range of proposed APOBEC-mediated DNA mutation targets in human cancer tissues and cell lines (Alexandrov et al. 2013; Petljak et al. 2019), especially in targets with dC/dT change (Nik-Zainal et al. 2012). This observation implies that APOBEC-mediated DNA and RNA editing frequency may each be functionally modified by AU enrichment in the flanking sequences surrounding modifiable bases. The base content in individual nucleotides surrounding the edited cytidine also exerted significant impact on editing frequency, particularly in a 10-nucleotide segment spanning the edited cytidine (Supplemental Table 1), accounting for 25% of the variance in editing frequency independent of the mooring sequence model. Our findings regarding individual nucleotides surrounding the edited cytidine are consistent with findings for both DNA and RNA editing targets, particularly in the setting of cancers (Backus and Smith 1992; Conticello 2012; Roberts et al. 2013; Saraconi et al. 2014; Gao et al. 2018; Arbab et al. 2020). Recent work examining the sequence-editing relationship of a large *in vitro* library of DNA targets edited by different synthetic cytidine base editor (CBE)s (Arbab et al. 2020) showed that the base content of a 6-nucleotide window spanning the edited cytidine explained 23-57% of the editing variance, in particular one or two nucleotides immediately 5’ of the edited nucleotide. That study also demonstrated that occurrence of T and C nucleotides at the position −1 increased, while a G nucleotide at that position decreased editing frequency (Arbab et al. 2020). However, in contrast to our findings, the presence of A at position −1 had either a negative or null effect on DNA editing activity (Arbab et al. 2020). This latter finding is consistent with the lower AU content observed in nucleotides adjacent to the edited cytidine in Apobec-1 DNA targets compared to the AU content in RNA targets. Our findings assign a greater importance of adjacent nucleotides in RNA editing frequency, similar to earlier reports that the five bases immediately 5’ of the edited cytidine in *Apob* mRNA exert a greater impact on editing activity compared to nucleotides further upstream of this segment (Backus and Smith 1991; Shah et al. 1991; Backus and Smith 1992). G/C fraction of a 6-nucleotide window spanning the edited cytidine in DNA targets is associated with editing activity of the synthetic CBEs (Arbab et al. 2020). Although we found significant associations of RNA editing with G/C fraction in segments surrounding the edited cytidine in univariate analyses, these associations were not retained in the final model. In contrast, the AU content of regulatory sequence motif B remained as an independent factor determining editing frequency in the final model.

The conserved 26-nucleotide sequence around the edited C forms a stem-loop secondary structure, where the editing site is in an octa-loop (Richardson et al. 1998) as predicted for the 55-nucleotide sequence of *ApoB* mRNA (Shah et al. 1991). This stem-loop structure is predicted to play an important role in recognition of the editing site by the editing factors (Bostrom et al. 1989; Davies et al. 1989; Driscoll et al. 1989; Chen et al. 1990). Mutations resulting in loss of base pairing in peripheral parts of the stem did not impact the editing frequency (Shah et al. 1991). Editing sites with the cytidine located in central parts (e.g. loop) exhibited higher editing frequencies than those with the edited cytidine located in peripheral parts (e.g. tail) and it is worth noting that the computer-based stem-loop structure was independently confirmed by NMR studies of a 31-nucleotide human *ApoB* mRNA (Maris et al. 2005). Those studies demonstrated that the location of the mooring sequence in the *ApoB* mRNA secondary structure plays a critical role in the RNA recognition by A1CF (Maris et al. 2005). In line with those findings, the current findings emphasize that the location of the mooring sequence in secondary structure of the target mRNA exerts significant independent impact on editing frequency. These predictions were confirmed in crystal structure studies of the carboxyl-terminal domain of APOBEC-1 and its interaction with cofactors and substrate RNA (Wolfe et al. 2020). Our conclusions regarding murine C-to-U editing frequency, such as mooring sequence, base content, and secondary structure appear consistent with a similar regulatory role among the smaller number of verified human targets. That being said, further study and expanded understanding of the range of C-to-U editing targets in human tissues will be needed as recently suggested (Destefanis et al. 2020), analogous to that for A-to-I editing (Bahn et al. 2012; Bazak et al. 2014).

We recognize that other factors likely contribute to the variance in RNA editing frequency not covered by our model. We did not consider the role of naturally occurring variants in APOBEC1, for example, which may be a relevant consideration since mutations in APOBEC family genes were shown to modify the editing activity of related hybrid DNA cytosine base editors (Arbab et al. 2020). Furthermore, genetic variants of *APOBEC1* in humans were associated with altered frequency of *GlyR* editing (Kankowski et al. 2017). Other factors not included in our approach included entropy-related features, tertiary structure of the mRNA target and other regulatory co-factors. Another limitation in the tissue-specific designation used to categorize editing frequency is that cell specific features of editing frequency may have been overlooked. For example, small intestinal and liver preparations are likely a blend of cell types (MacParland et al. 2018; Elmentaite et al. 2020) and tumor tissues are highly heterogeneous in cellular composition (Barker et al. 2009). The current findings provide a platform for future approaches to resolve these questions.

## MATERIALS AND METHODS

### Search strategy

A comprehensive literature review from 1987 (when *ApoB* RNA editing was first reported (Chen et al. 1987; Powell et al. 1987)) to November 2020, using studies published in English reporting C-to-U mRNA editing frequencies of individual or transcriptome-wide target genes. Databases searched included Medline, Scopus, Web of Science, Google Scholar, and ProQuest (for thesis). The references of full texts retrieved were also scrutinized for additional papers not indexed in the initial search.

### Study selection

Primary records (N=528) were screened for relevance and *in vivo* studies reporting editing frequencies of individual or transcriptome-wide APOBEC1-dependent C-to-U mRNA targets selected, using a threshold of 10% editing frequency. For analyses based on RNA sequence information, only targets with available sequence information or chromosomal location for the edited cytidine were included. Exclusion criteria included: studies that reported C-to-U mRNA editing frequencies of target genes in other species, studies reporting editing frequencies of target genes in animal models overexpressing APOBEC1, exclusively *in vitro* studies, and conference abstracts.

### Human targets

We included studies reporting human C-to-U mRNA targets (Chen et al. 1987; Powell et al. 1987; Skuse et al. 1996; Mukhopadhyay et al. 2002; Grohmann et al. 2010; Schaefermeier and Heinze 2017). We also included work describing APOBEC1-mediated mutagenesis in human breast cancer (Nik-Zainal et al. 2012).

### Data extraction

Two reviewers (SS and VB) conducted the extraction process independently and discrepancies were addressed upon consensus and input from a third reviewer (NOD). The parameters were categorized as follows: *General parameters*: Gene name (RNA target), chromosomal and strand location of the edited cytidine, tissue site, editing frequency determined by RNA-seq or Sanger sequencing as illustrated for *ApoB* (Figure 1A). Editing frequency was highly correlated by both approaches (*r*=0.8 *P*<0.0001), and where both methodologies were available we used RNA-seq. We also defined relative dominance of editing co-factors (A1CF-dominant, RBM47-dominant, or co-dominant), relative mRNA expression (edited gene vs unedited gene) by RNA-seq or quantitative RT-PCR, and abundance of corresponding protein (edited gene vs unedited gene) by western blotting or proteomic comparison. Co-factor dominancy was determined based on the relative contribution of each co-factor to editing frequency. In each editing site, editing frequencies in mouse tissues deficient in *A1cf* or *Rbm47* were compared to that of wild-type mice. The relative contribution of each co-factor was calculated by subtracting the editing frequency for each target in *A1cf* or *Rbm47* knockout tissue from the total editing frequency in wild-type control. Editing sites with <20% difference between contributions of RBM47 and A1CF were considered co-dominant. Sites with ≥20% difference were considered either RBM47-or A1CF-dominant, depending on the co-factor with higher contribution (Blanc et al. 2019).

#### Sequence-related parameters

A sequence spanning 10 nucleotides upstream and 30 nucleotides downstream of the edited cytidine was extracted for each C-to-U mRNA editing site. These sequences were extracted either directly from the full-text or using online UCSC Genome Browser on Mouse (NCBI37/mm9) and Human (Grch38/hg38) (https://genome.ucsc.edu/cgi-bin/hgGateway). Using the mooring sequence model (Backus and Smith 1992), three *cis*-acting elements were considered for each site. These elements included 1) a 10-nucleotide segment immediately upstream of the edited cytidine as “regulatory sequence”; 2) a 10-nucleotide segment downstream of the edited cytidine with complete or partial consensus with the canonical “mooring sequence” of *ApoB* mRNA; 3) the sequence between the edited cytidine and the 5’ end of the mooring sequence, referred to as “spacer”. We used an unbiased approach to identify potential mooring sequences by taking the nearest segment to the edited cytidine with lowest number of mismatch(es) compared to the canonical mooring sequence of *ApoB* RNA. For each of the three segments, we investigated the number of mismatches compared to the corresponding segment of *ApoB* gene (Blanc et al. 2014), as well as length of spacer, the abundance of A and U nucleotides (AU content) and the G to C abundance ratio (G/C fraction (Arbab et al. 2020)). We also calculated relative abundance of A, G, C, and U individually across a region 10 nucleotides upstream and 20 nucleotides downstream of the edited cytidine across all editing sites. For comparison, we examined the base content of a sequence spanning 10 nucleotides upstream and downstream of mutated deoxycytidine for over 6000 proposed C to X (T, A, and G) DNA mutation targets of APOBEC family in human breast cancer (Nik-Zainal et al. 2012) along with relative deoxynucleotide distribution in proximity to the edited site.

#### Secondary structure parameters

We used RNA-structure (Reuter and Mathews 2010) and Mfold (Zuker 2003) to determine the secondary structure of an RNA cassette consisting of regulatory sequence, edited cytidine, spacer, and mooring sequence. Secondary structures similar to that of the cassette for *ApoB* chr12: 8014860 consisting of one loop and stem (with or without unassigned nucleotides with ≤4 unpaired bases inside the stem) as the main stem-loop with or without free tail(s) in one or both ends of the stem were considered as canonical. Two other types of secondary structure were considered as non-canonical structures (Figure 1B), with ≥2 loops located either at ends of the stem or inside the stem. Loops inside the stem were circular open structures with ≥5 unpaired bases. Editing sites with canonical structure were further categorized into three subgroups based on location of the edited cytidine: specifically (C_loop_), stem (C_stem_), or tail (C_tail_). In addition to overall secondary structure, we considered location of the edited cytidine, location of mooring sequence, symmetry of the free tails, and proportion of the nucleotides in the target cassette that constitute the main stem-loop. This proportion is 1.0 in the case of *ApoB* chr12: 8014860 where all the bases are part of the main stem-loop structure. Symmetry was defined based on existence of free tails in both ends of the RNA strand.

### Statistical methodology

Continuous variables are reported as means ± SD with relative proportions for binary and categorical variables. *T-test* and *ANOVA* tests were used to compare continuous parameters of interest between two or more than two groups, respectively. *Chi-squared* testing was used to compare binary or categorical variables among different groups. Pearson *r* testing was used to investigate correlation of two continuous variables. We used linear regression analyses to develop the final model of independent factors that correlate with editing frequency. We used the Hosmer and Lemeshow approach for model building (Hosmer Jr et al. 2013) to fit the multivariable regression model. In brief, we first used bivariate and/or simple regression analyses with P value of 0.2 as the cut-off point to screen the variables and detect primary candidates for the multivariable model. Subsequently, we fitted the primary multivariable model using candidate variables from the screening phase. A backward elimination method was employed to reach the final multivariable model. Parameters with P values <0.05 or those that added to the model fitness were retained. Next, the eliminated parameters were added back individually to the final model to determine their impact. Plausible interaction terms between final determinants were also checked. The final model was screened for collinearity. We used the same approach to develop a multinomial logistic regression model to identify factors that were independently associated with co-factor dominance in RNA editing sites. Squared R and pseudo squared R were used to estimate the proportion of variance in responder parameter that could be explained by multivariable linear regression and multinomial logistic regression models, respectively. The same screening and retaining methods were used to investigate association of base content in a sequence 10 nucleotides upstream and 20 nucleotides downstream of the edited cytidine, with editing frequency. However, after determining the nucleotides that were retained in final regression model, a proxy parameter named “base content score” was calculated for each editing site based on the β coefficient values retrieved for individual nucleotides in the model. This parameter was used in the final model as representative variable for base content of the aforementioned sequence in each editing site.

## ACKNOWLEDGMENTS

This work was supported by grants from the National Institutes of Health grants DK-119437, DK-112378, Washington University Digestive Diseases Research Core Center P30 DK-52574 (to NOD)

## SUPPLEMENTAL FIGURE LEGENDS

**Supplemental Figure 1. Chromosomal distribution of murine APOBEC1-mediated C-to-U mRNA editing sites**. The black curve corresponds to left Y-axis and represents average editing frequencies of editing sites related to each chromosome. The blue curve corresponds to right Y axis and represents number of editing sites related to each chromosome.

**Supplemental Figure 2. Association of editing frequency with characteristics of regulatory sequence in murine APOBEC1-mediated C-to-U mRNA editing sites**. A-C. Association of editing frequency with number of mismatches and AU content (%). D-F Association of editing frequency with different regulatory sequence motifs. Mismatches were determined in comparison to the same regulatory sequence motif in *ApoB* mRNA (as reference).

**Supplemental Figure 3. Association of editing frequency with characteristics of downstream sequence in murine APOBEC1-mediated C-to-U mRNA editing sites**. A. Association of editing frequency with spacer length. B. Association of editing frequency with spacer AU content (%). C-F. Association of editing frequency with and AU content of successive segments downstream of the edited cytidine.

**Supplemental Figure 4. Association of editing frequency with secondary structure-related characteristics in C-to-U mRNA editing sites**. A: distribution of edited cytidine location in secondary structure regardless of the overall secondary structure. B: association of editing frequency with edited cytidine location in secondary structure. C: distribution of free tail orientation in editing sites. D: association of editing frequency with free tail orientation in editing sites. E: association of editing frequency with 3’ free tail length. * *P*<.05; *** *P*<.0001. *r*: Pearson correlation coefficient.

**Supplemental Figure 5. Association of secondary structure-related characteristics with dominant co-factor pattern in APOBEC1-mediated C-to-U mRNA editing sites**. A. Distribution of mooring sequence location presented in the context of different dominant co-factor patterns. B. Distribution of free tail orientation in secondary structure among editing sites, presented in the context of different dominant co-factor patterns.

**Supplemental table 1.**
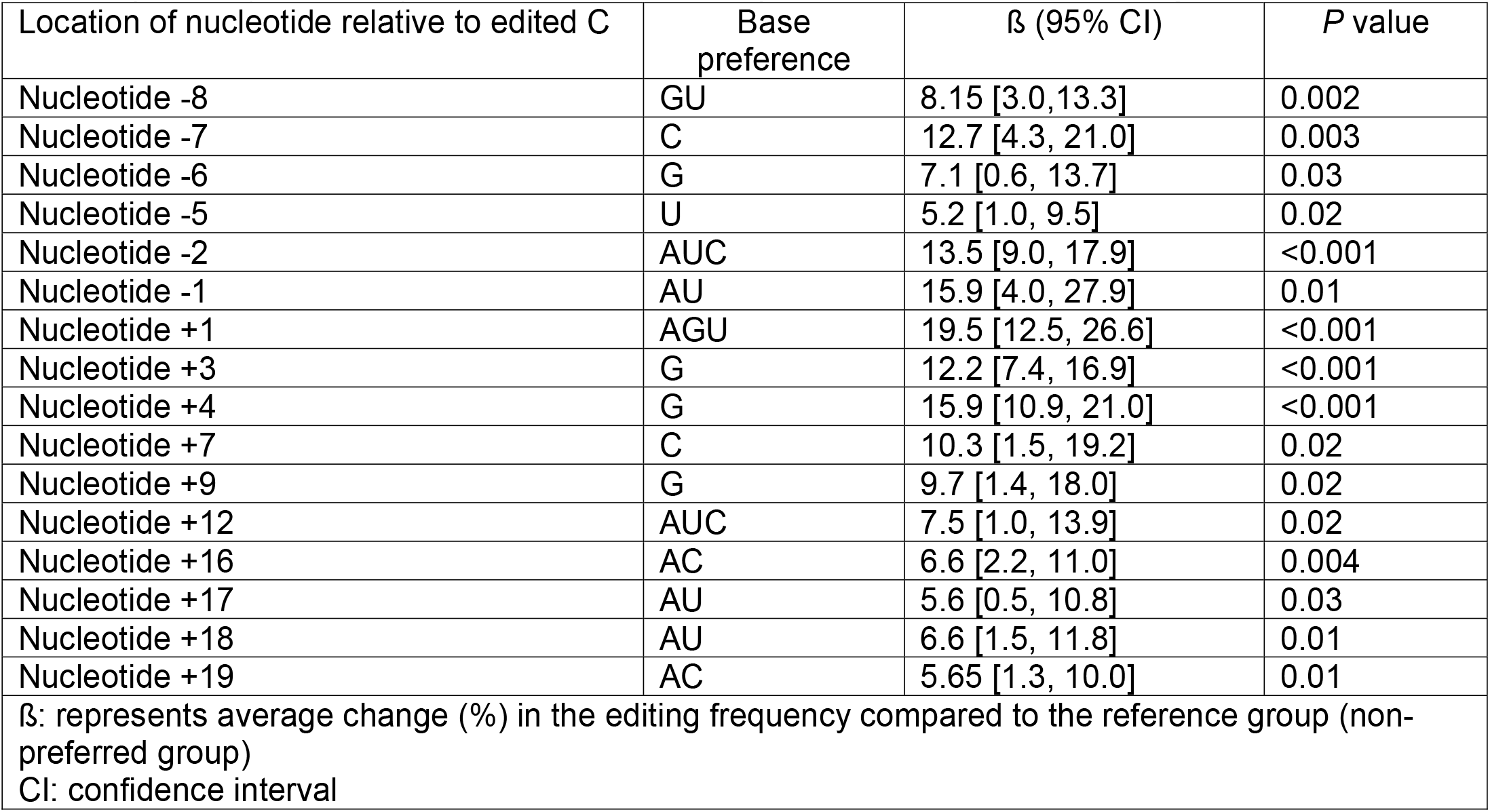
Multivariable linear regression model for individual nucleotides surrounding edited cytosine (−10 to +20) in mouse APOBEC1-dependent C-to-U mRNA editing sites.

**Supplemental table 2.**
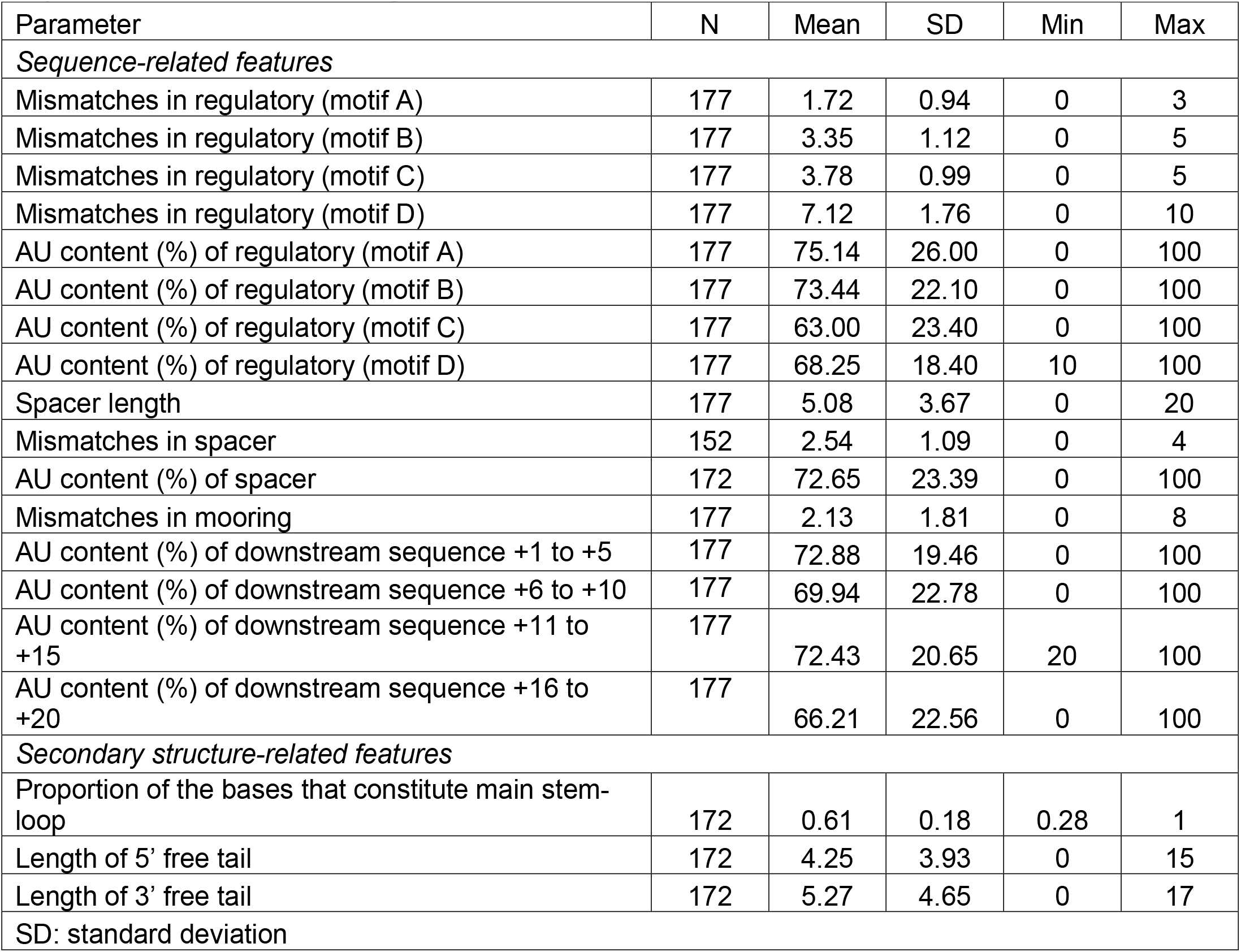
Descriptive data of regulatory-spacer-mooring cassette in mouse APOBEC1-dependent C-to-U mRNA editing sites.

**Supplemental table 3.**
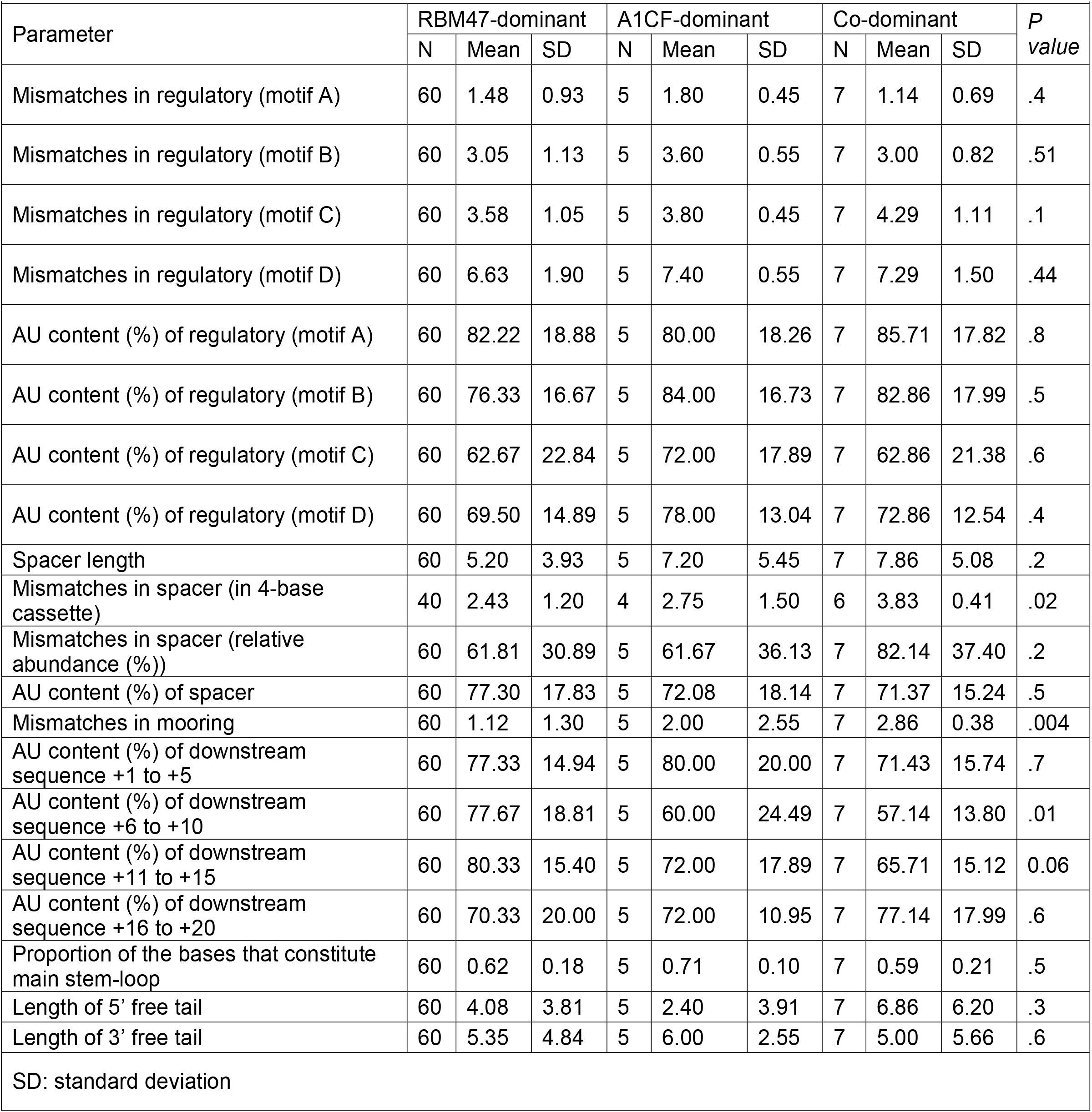
Comparing three subgroups of mouse APOBEC1-dependent C-to-U mRNA editing sites based on co-factor dominance.

**Supplemental Table 4.**
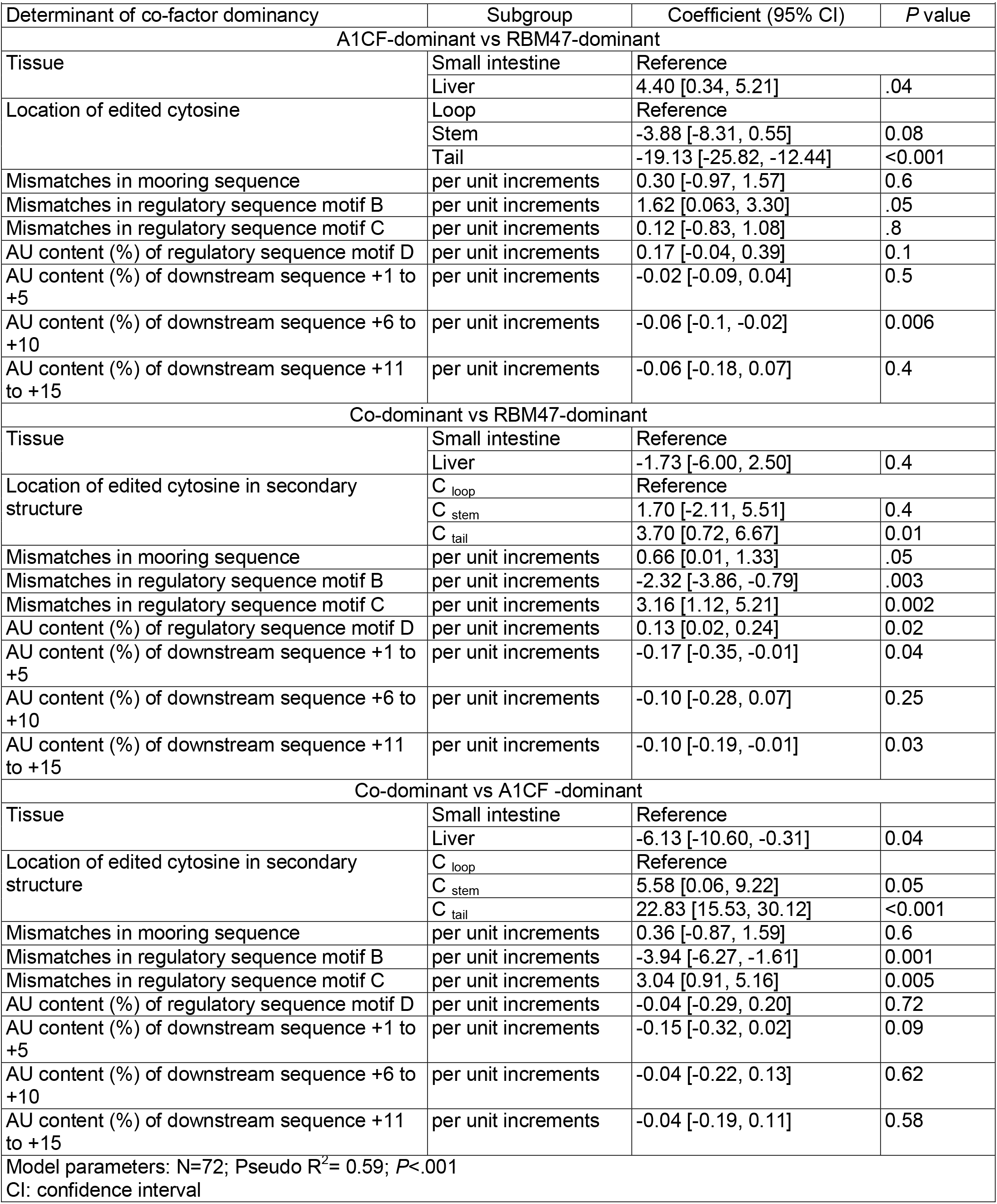
Multinomial logistic regression model for determinant factors of co-factor dominancy in mouse APOBEC1-dependent C-to-U mRNA editing sites.

